# Characterisation of novel bacteriophages against the cattle pathogen *Moraxella bovis*

**DOI:** 10.1101/2025.11.24.690234

**Authors:** Hannah R Sampson, Malgorzata Wegrzyn, Theo Josephs, Nzubechukwu I Ugokwe, Andrew Kinsella, Anisha M Thanki, Deepinder K Kalra, Alexane Roux, Hannah L Patrick, Benjamin MC Swift, Gregory Firth, Raj Odedra, Andrew D Millard, Martha RJ Clokie

**Affiliations:** Becky Mayer Centre for Phage Research, Division of Microbiology and Infection, University of Leicester, UK; Carus Animal Health, Surrey, UK

## Abstract

**Background:** Infectious bovine keratoconjunctivitis is the most important cattle ocular disease worldwide. The infection is primarily caused by *Moraxella bovis* and is a highly contagious disease that significantly affects cattle welfare. Currently, antibiotic medication is the primary treatment for infectious bovine keratoconjunctivitis. However, with rising concerns over antibiotic resistance, we propose developing a more targeted therapeutic strategy using bacteriophages (phages).

**Materials and Methods:** We have isolated the first known *Moraxella bovis* phages, characterised them according to their genome sequence, local virulence index and with transmission electron microscopy. The host ranges were assessed using 41 clinical *M. bovis* strains isolated from infected cows.

**Results:** Four phages were isolated and characterised. Comparative analysis identified a high degree of genomic similarity between the phages MB15, MB16, MB26 and MB43. MB43 was the most distinct, with the smallest host range phenotype.

**Conclusions:** The isolated phages show therapeutic potential for further development against *Moraxella* infections.

## Introduction

*Moraxella bovis* is the primary etiological agent of infectious bovine keratoconjunctivitis (IBK), the most common ocular disease of cattle worldwide^1–3^. IBK, also known as pinkeye or New Forest Eye, is a highly contagious disease that has a considerable negative impact on cattle welfare. Risk factors for IBK include exposure to UV radiation from sunlight, dry or dusty conditions and direct trauma to the eye from irritants such as pollen or grasses^4^. IBK progression is marked by worsening corneal opacity and ulceration, in severe cases, IBK can lead to blindness^3,5^.

Previous studies investigating the bovine bacterial ocular microbiome have identified bacteria from the genera *Mycoplasma* and *Moraxella*, family *Weeksellacea* (genus *Bergeyella*) and family *Pasteurellaceae* as the most common taxa^6–8^. The prevalence of *Moraxella* and *Mycoplasma* correlates with IBK, and other pathogens may worsen disease severity^4,8,9^. *Moraxella bovis* has long been linked with IBK^3,10^, but other species, such as *Moraxella bovoculi* are often associated with infection sites and may play a role in disease, but this has not been empirically shown^11–13^.

The nature of *M. bovis* pathogenicity in IBK has been characterised with two distinct stages. Initially, *M. bovis* adheres to the corneal epithelium with pili^14–18^. After adherence, *M. bovis* secretes haemolytic cytotoxins that subsequently lyse the corneal epithelial cells^19,20^. An assessment of *M. bovis* genomes identified two major genotypes, with genotype 2 that carry more virulence genes and prophages than genotype 1^21^. *M. bovis* genotype is important to consider in IBK, as virulence factors vary among isolates, which affect their fitness and roles in disease.

Antibiotics, including oxytetracycline, cloxacillin and tulathromycin, are the primary treatment for IBK^4,22–24^, with attempts made for vaccination but these have lacked efficacy^25,26^. *M. bovis* antimicrobial resistance patterns overall vary by isolate^22,23^, but resistance pattern trends have also been observed in *M. bovoculi* isolates from the same geographical origin^23^. In addition to the negative impacts on cattle welfare, IBK presents a significant economic impact to the cattle industry that can be attributed to corneal ulceration, a reduction in cattle weight gain at weaning^1^ and the associated additional costs of treatment such as repeated antibiotic applications.

Therefore, we propose an alternative IBK treatment using bacteriophages (phages) that specifically target *M. bovis* to treat this difficult and debilitating disease. Phage-like particles have previously been observed from *M. bovis* pilin preparations, yet *M. bovis* phages were not isolated^27^.

In this study we isolated and purified four *Moraxella bovis* phages from a cow nasal swab. Phages were characterised using transmission electron microscopy, their local virulence index was calculated, and host range was assessed against a library of clinical *M. bovis* strains. Phage genomes were assembled, annotated, compared and taxonomically identified as a new genus within the *Straboviridae* family, named “Nasusvirus vacca*”* (named after the location the phages were isolated, from Latin ‘nasus’ nose and ‘vacca’ cow). This research introduces these novel phages and provides a solid foundation for future studies into the therapeutic potential of these phages as an IBK treatment for *Moraxella* infectious disease.

## Methods

### Bacterial culture conditions

The *Moraxella bovis* reference strain (NCTC 9426) was used in this study as this strain has been characterised and sequenced^28–31^. The *M. bovis* clinical isolates (Table S1) were provided by Orhan Sahin (Iowa State University). *M. bovis* strains were stored at -80 °C in growth medium supplemented with 20 % glycerol and bacteria were cultivated using blood agar (Oxoid) supplemented with 5 % (v/v) horse blood (Oxoid) and incubated at 37 °C overnight. Bacterial cultures were grown in brain heart infusion broth (BHI, Oxoid) supplemented with 1 mM CaCl_2_ (BHI+) and incubated at 200 rpm, 37 °C overnight (16 to 18 h). Fresh BHI+ was inoculated with bacterial overnight culture to an optical density (OD_600nm_) of 0.1 and incubated at 200 rpm, 37 °C until mid-exponential growth phase (∼ 0.7 OD_600nm_). Unless otherwise stated, bacterial lawns were prepared by mixing 750 µL of bacterial culture with 1.25 mL of molten 0.5 % BHI agar, supplemented with 1 mM CaCl_2_ (Fisher) and poured onto BHI agar plates.

### Bacteriophage isolation

For phage isolation, a swab (Heinz Herenz LMS-Swab, Amies) was used to sample the nasal passages of a cattle cadaver. The swab sample was initially incubated with 3 mL of SM buffer (100 mM NaCl, 8 mM MgSO4 · 7H2O, and 50 mM Tris-HCI) on a rotator overnight. The sample was subsequently 0.45 µm filtered and 600 µL was used for enrichment in 4 mL of fresh BHI+ with the addition of 50 µL *M. bovis* NCTC 9426 overnight culture. The enrichment was incubated at 200 rpm, 37 °C overnight. Samples were centrifuged at 3095 RCF for 10 min and the supernatant was 0.22 µm filtered. The enrichment was carried out on the filtrate a second time and the resulting filtrate was tested for the presence of phages using the agar overlay method^32^. Plates were incubated at 37 °C overnight and clear plaque formation was observed.

### Bacteriophage purification and storage

Phages were purified by three rounds of plaque pick purification^32^. As phage were unstable at 4 °C (Figure S1), phages were kept at –80 °C for long term storage. To create stocks, phage was mixed with mid exponential phase *M. bovis* NCTC 9426 and incubated for 15-20 min at room temperature, prior to the addition of glycerol at final concentration of 20 % (v/v). To revive phages from –80 °C storage, an aliquot of phage infected cells was defrosted at room temperature, added to 5 mL of fresh BHI+ and incubated overnight at 200 rpm, 37 °C. Samples were centrifuged at 3095 RCF for 10 min and supernatant was 0.22 µm filtered.

### Assessing bacteriophage morphology with transmission electron microscopy (TEM)

Phage lysates were purified with ammonium acetate and were stained with 1 % x/v uranyl acetate following a previously established method^33^. The stain was left to stand for 10 s and excess stain was removed by blotting. The samples were imaged using JEOL JEM-1400 TEM with an accelerating voltage of 120 kV. Digital images were captured using an EMSIS Xarosa 20MP digital camera with Radius software and analysed with ImageJ^34^.

### Bacteriophage titre

Phage titres were determined using the agar overlay method^32^. A tenfold dilution series of phage lysate was made using SM buffer, each dilution was plated in triplicate on the agar overlay and incubated at 37 °C overnight. Plaques were enumerated, and plaque forming units per millilitre (PFU/mL) were calculated.

### Host range

Host range of isolated phages were determined against a library of clinical *M. bovis* isolates (Table S1) using an agar overlay spot assay^32^. Briefly, 750 µL of 18 h overnight bacterial culture was mixed with 2.5 mL of molten 0.5 % BHI agar, supplemented with 1 mM CaCl_2_ and 1 mM MgSO_4_, and poured onto BHI agar plates. Phage lysates were serially diluted and spotted onto solidified bacterial lawns; plates were incubated at 37 °C for 18-24 h. The presence of plaques and lysis zones was recorded and categorised.

### One-Step

A one-step experiment was carried out based on a previously established method^33,35^. Briefly, an overnight culture was diluted 1:100 and grown to 0.4 OD_600nm_. Phages were added to a final multiplicity of infection (MOI) of ∼0.01 and allowed to absorb for 1 min at 37 °C shaking at 300 rpm. Free phages were removed by centrifugation at 13000 x g for 1 min, the supernatant was removed, and the bacterial pellet resuspended in fresh BHI+. Samples were taken at regular intervals for 90 min (every 15 min up to 30 min and then every 5 min up to 50 min with 15 min intervals thereafter until the end point). Phage titre was determined using a double agar overlay plaque assay.

### Adsorption assay

An adsorption assay was carried out based on a previously established method^33^. Briefly, overnight culture was diluted 1:100 and grown to 0.4 OD_600nm_. MB26 phages were added to a final MOI of ∼0.01 and thoroughly mixed. At each timepoint, a sample was taken, centrifuged at 13000 x g for 1 min and the supernatant was transferred to a fresh tube. Samples were taken at 2 min intervals for 28 min and free phage titre was determined using a double agar overlay plaque assay.

#### Local Virulence Index

A local virulence index (VI) assay was carried out for three MOI using a modified method based on previous work^36^. Briefly, overnight culture was diluted 1:100 and grown to 0.1 OD_600nm_, equating to ∼10^7^ of bacterial cells/mL. Bacterial cultures were added to triplicate wells and infected with phages in a 1:1 ratio at an MOI of 0.01, 0.001 or 0.0001 to a final volume to 200 µl. Plates were incubated at 37 °C for 24 h and OD_600nm_ was measured with BMG labtech Spectrostar Nano Plate Reader. Optical density readings were compared to bacteria and media controls, and the local VI was calculated from 500 min of growth of three biological repeats.

The local VI was calculated (1-(φi/φc)) in GraphPad Prism (v10.4.0 for Windows, GraphPad Software, USA) using the area under the curve of the phage of interest (φi) at a specific MOI divided by area under the curve of the control culture (φc). Data were visualised in R studio^37^.

### DNA extraction, sequencing, genome assembly and annotation

Phage DNA was extracted using a modified version of a previously described phenol extraction method^33,38^. Briefly, prior to extraction, phage lysate was treated with DNase I (1 mg/mL) for 1 h at 37 °C. An equal volume of phenol:choroform:isoamyl alcohol was added to the sample, mixed, and left for 2 min, prior to centrifugation at 13000 x g for 10 min. The aqueous layer was extracted, and the process repeated three times. The aqueous layer was mixed with 1/10 volume of 3 M of sodium acetate, and two volumes of 100% ethanol and DNA was precipitated at –20 °C. DNA pellets were washed twice with ethanol. The supernatant was discarded, DNA pellets were air dried, then resuspended in 50 µl of nuclease free water.

DNA was sequenced by using llumina NovaSeq X Plus NexteraXT at SeqCenter, using tagmentation library preparation, producing paired sequences with a length of 150 bp. For genome assembly, reads were filtered and trimmed with BBMap (v39.08) (bbduk paired read settings: ktrim=r k=23 mink=11 hdist=1 tpe tbo), Trim Galore (v 0.6.10) and coverage was normalised with bbnorm (target=150). Genomes were assembled using SPades^39^ v4.0.0 (only assembler), orientated with DNAppler v1.1.0, polished using pilon v1.24 and annotated with Pharokka^40^ v1.7.3 run with threads (8) and PHANOTATE^41^ for gene prediction. Taxonomy was determined using taxMyPhage^42^. Genomes were compared using dnadiff^43,44^ (mummer4 v4.0.1), Roary^45^ (v3.13.0) and visualised with clinker^46^. Genome information is detailed in Table S2 and sequences are available under BioProject (PRJEB81897).

Phylogenetic analysis was carried out with ViPTree^47^ (v1.1.2), with default parameters using a database of genomes to represent all genera within the order *Pantevenvirales* as defined in the VMR_MSL_V40.1 . The alignment was visualised in iTOL^48^.

## Results and Discussion

Four phages MB15, MB16, MB26 and MB43, were isolated from a single cattle nasal swab that was used to sample an individual cadaver. Phages were isolated following two rounds of enrichment with the *M. bovis* reference *s*train NCTC 9426. All four phages exhibited similar myovirus morphologies and sizes (Figure 1, Table S3), featuring a large, elongated head, contractile tail and complex base plate.

**Figure 1.**
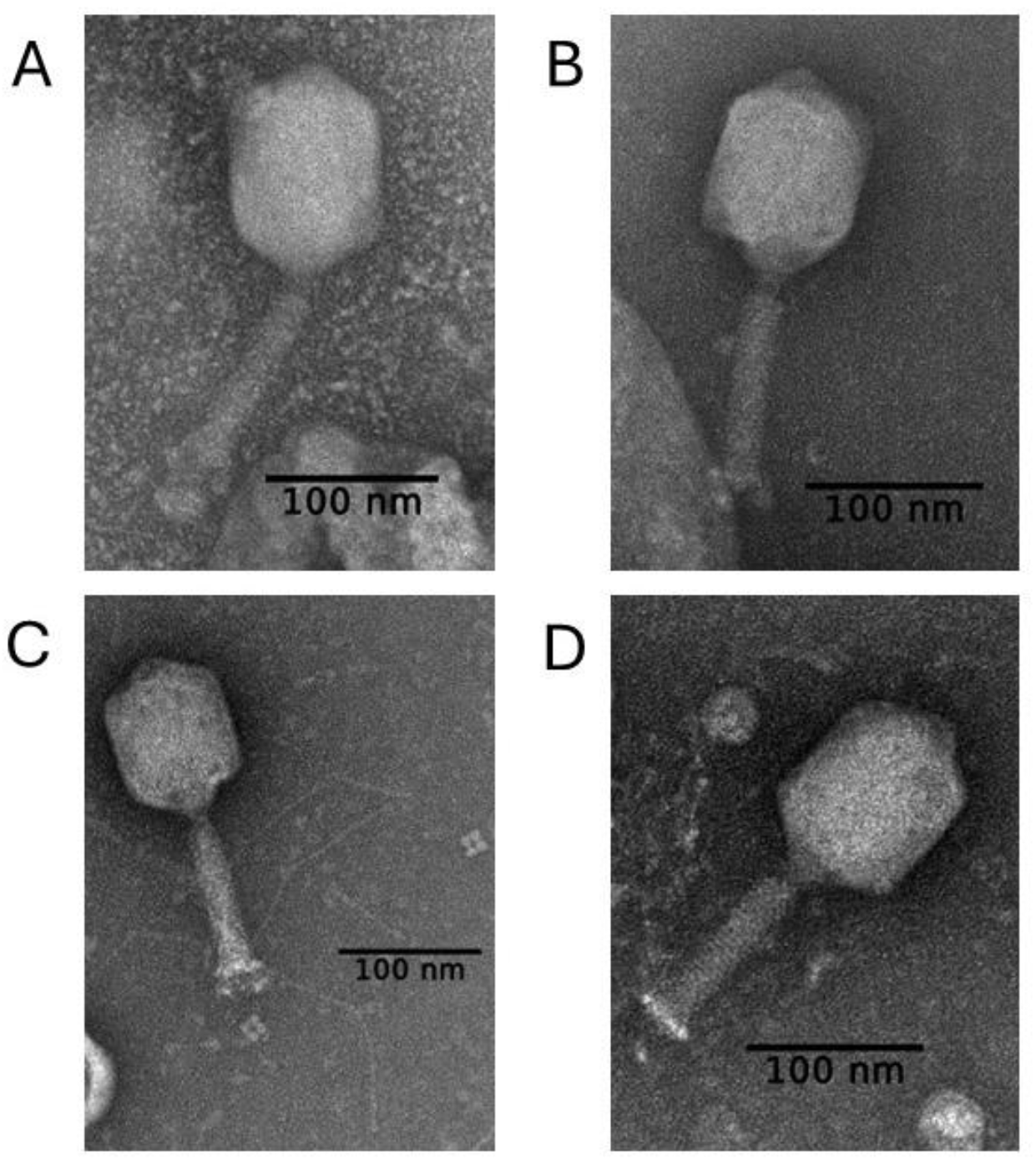
Morphologies of four isolated *M. bovis* phages (A) MB15, (B) MB16, (C) MB26 and (D) MB43. Phages were isolated from cattle nasal swab samples following enrichment with *M. bovis* NCTC 9426. ImageJ was used to measure phage dimensions of three virions. Measurements are detailed in Table S3.

The local VI for phages at three different MOI was calculated against the *M. bovis* NCTC 9426 host (Figure 2A, Table S4). The mean local VI values increased with higher MOI, each MOI level differed significantly from the others (p <0.05, Table S5, Table S6). MB16 was significantly different to all other phages at MOI 0.01, with the highest local VI of 0.77 (Figure 2A, Table S7). At MOI of 0.0001, MB26 had a significantly higher local VI then all other phages, and MB15 had a significantly lower local VI than all other phages, no other significant differences were found between phages (Table S7).

**Figure 2.**
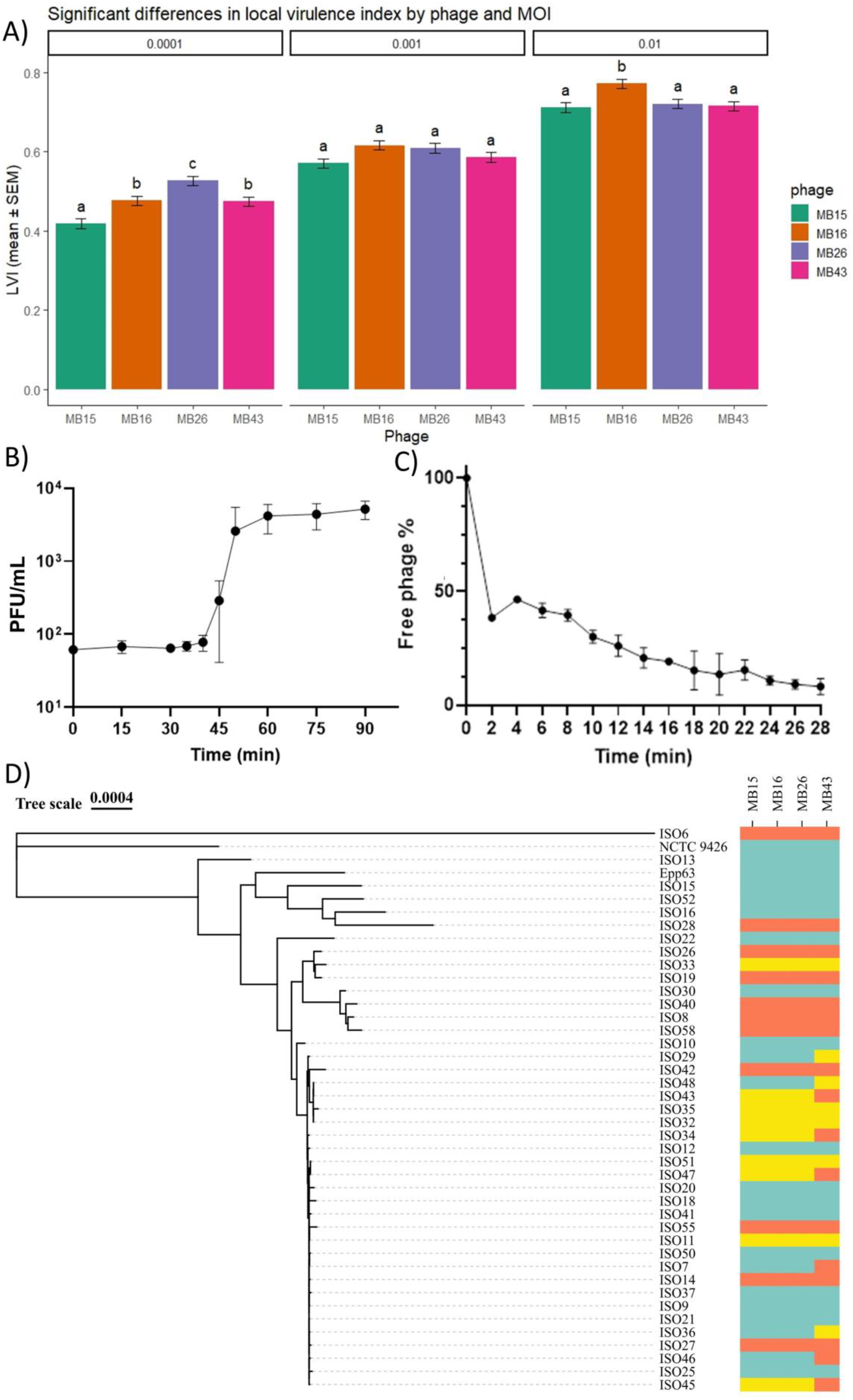
A) Local virulence index (LVI) of phages MB15, MB16, MB26 and MB43 at three different MOI (0.01, 0.001 and 0.0001) against host strain *M. bovis* NCTC 9426. Unique letters represent significant differences between phages at the 0.05 level. Bars represent standard error of the mean, n=3. B) One-Step growth curve of MB26 propagated in isolation host *M. bovis* NCTC 9426 at MOI 0.01. The mean and standard deviation are plotted, n=3. C) Adsorption rate of phage MB26 in isolation host *M. bovis* NCTC 9426 at MOI 0.01. D) Host range of phages MB15, MB16, MB26 and MB43 against clinical *M. bovis* strains, phage isolation strain *M. bovis* NCTC 9426 and *M. bovis* reference strain Epp-63. Bacterial phylogeny was created using REALPHY^51,52^, phage effectivity is represented by colour, where teal: visible plaques, yellow: clearing effect or inconsistent visible plaques and orange: no visible effect on bacterial lawn.

MB26 was further characterised with a One-Step growth curve (Figure 2B) and adsorption assay (Figure 2C) with *M. bovis* NCTC 9426 isolation host (bacterial growth curve Figure S2). The burst size of MB26, was 71.25 PFU/mL. An adsorption assay determined that after 2 min, >50% of phage MB26 was absorbed, and at 26 min, 90% was adsorbed into host *M. bovis* (Figure 2C).

Host range was tested against a library of clinical *M. bovis* isolates. Phages infected 21 of 41 clinical *M. bovis* strains (Figure 2D, Table S8) and all phages infected *M. bovis* reference strain Epp-63^49^. Phage MB43 had the narrowest host range, infecting 16 out of 41 clinical strains. In general phages had largely similar host ranges, with variation in plaquing efficiency (Table S8). Interestingly, the use of mid-exponential bacterial cultures for lawns, gave slightly different host range results (e.g. ISO10, ISO11 and ISO12; Table S9) compared to overnight lawns (Table S8). This demonstrates that phage infectivity is partially dependent on bacterial growth phase, perhaps linked to *M. bovis* phase variation^16,21,50^.

Genome analysis of MB15, MB16, MB26 and MB43 revealed that phage genomes ranged from 123294 bp to 123307 bp, with a mol GC content of ∼36 % (Table S2). MB15 contained 161 predicted genes, MB43 163 genes and MB26 and MB16 both contain 164 genes. All phages had 9 tRNA genes. Analysis with PhageLeads^53^ identified the phages did not contain any known genes associated with lysogeny, increasing bacterial virulence or conferring antimicrobial resistance (Figure S3). The sequencing approach used in this study did not allow determination of genomic termini or detection of nucleotide modifications; therefore, these features were not analysed.

Analysis with taxMyPhage demonstrated the phages do not fall within any known genus or species, and all four phages are the same species, given the >95% similarity to each other^54^. Further phylogenetic analysis with VipTree, placed the phages within the family *Straboviridae* (Figure 3A), in a sister clade to phages in the genus *Lasallevirus*. Thus, we proposed these phages form a new genus and species within the family *Straboviridae*, named “Nasusvirus vacca” (named after the location from which the phages were isolated, from Latin ‘nasus’ nose and ‘vacca’ cow).

**Figure 3:**
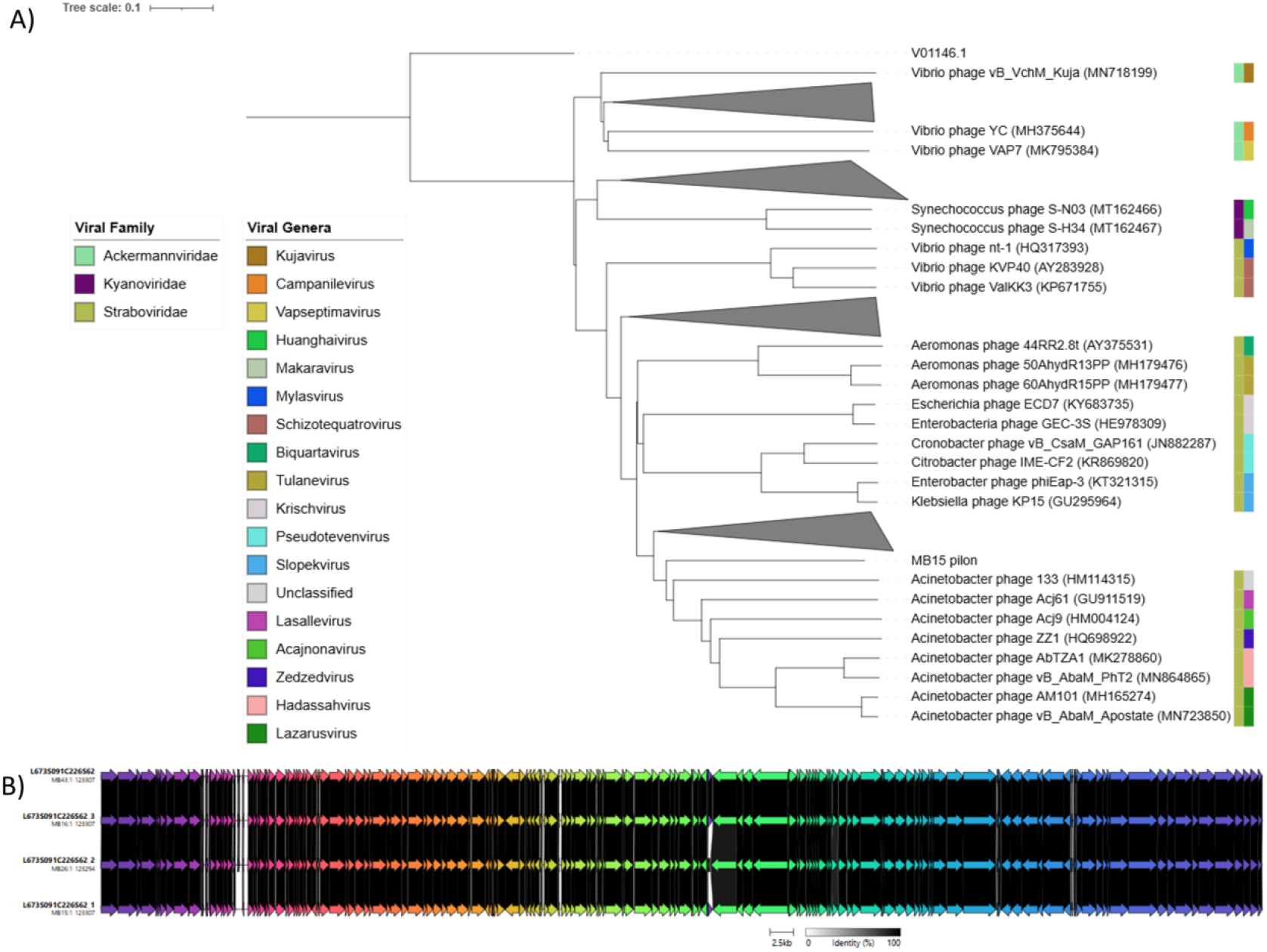
A) Phylogenetic tree of *Moraxella bovis* phage MB15. Colours represent viral family and viral genera as denoted by the legends. MB15 was classified in the *Straboviridae* family and the closest relatives to MB15 was identified as *Acinetobacter* phage 133 (HM114315) and *Acinetobacter* phage Acj61 (GU911519) in the *Lasallevirus* genus. Clades of similar individuals in outgroups are collapsed (in grey) for viewing clarity. B) Genome alignment of *M. bovis* bacteriophages MB15, MB16, MB26 and MB43 showing regions of gene similarities in black. Clinker was used for genome alignments, the best links identified by the default parameters are displayed.

Comparative analysis identified a high degree of genomic similarity (Figure 3B) with eight gene changes and 35 single nucleotide polymorphism (SNP) differences. All phages had unique SNPs (Table 1), four genes were unique to one phage and a further four were present in two or three phages suggesting they are part of their accessory genome (Table S10, Table S11). As most gene changes were listed as hypothetical genes, their functional roles remain uncertain. MB26 has deletions in long tail fiber protein distal subunit HRSMB15_0101 in comparison to the other phages (Tables S12-S17) which account for the decreased genome length and may confer changes in host preferences, availability and the infection process^55^.

**Table 1:** Unique SNP differences in MB15, MB16, MB26 and MB43.

| Phage | SNP | Comp NT | Comp genome | Gene ID | GeneName | Change | AA change |
| --- | --- | --- | --- | --- | --- | --- | --- |
| MB15 | A | G | MB16 | - | - | intergenic | - |
| MB15 | G | T | MB16 | - | - | intergenic | - |
| MB15 | C | T | MB16 | HRSMB15_0133 | Alt-like RNA polymerase ADP-ribosyltransferase | nonsynonymous | L94358F |
| MB15 | C | T | MB16 | HRSMB15_0145 | baseplate wedge subunit | synonymous | F103688F |
| MB15 | C | A | MB16 | HRSMB15_0151 | baseplate wedge subunit | nonsynonymous | N109471K |
| MB16 | G | A | MB26 | - | - | intergenic | - |
| MB16 | G | T | MB26 | HRSMB16_0090 | RNA ligase | nonsynonymous | V56534L |
| MB16 | T | G | MB26 | HRSMB16_0136 | Alt-like RNA polymerase ADP-ribosyltransferase | nonsynonymous | V94353G |
| MB16 | G | T | MB26 | HRSMB16_0136 | Alt-like RNA polymerase ADP-ribosyltransferase | nonsynonymous | L94360F |
| MB16 | G | T | MB26 | HRSMB16_0136 | Alt-like RNA polymerase ADP-ribosyltransferase | nonsynonymous | E94363D |
| MB16 | G | T | MB26 | HRSMB16_0136 | Alt-like RNA polymerase ADP-ribosyltransferase | synonymous | S94369S |
| MB16 | C | A | MB26 | HRSMB16_0154 | baseplate wedge subunit | nonsynonymous | L109460I |
| MB16 | G | A | MB43 | HRSMB16_0023 | hypothetical protein | synonymous | A14614A |
| MB26 | G | T | MB43 | HRSMB26_0023 | hypothetical protein | nonsynonymous | R14617S |
| MB26 | G | T | MB43 | HRSMB26_0090 | RNA ligase | nonsynonymous | L56533F |
| MB26 | G | T | MB43 | HRSMB26_0096 | DNA topoisomerase II | nonsynonymous | G62372V |
| MB26 | T | C | MB43 | HRSMB26<br>_0104 | long tail fiber<br>protein distal<br>subunit | synonymous | K66614K |
| MB26 | G | T | MB43 | HRSMB26<br>_0155 | baseplate<br>wedge subunit | nonsynonymous | C112951F |
| MB43 | G | T | MB43 | HRSMB26<br>_0023 | hypothetical<br>protein | nonsynonymous | G14607V |
| MB43 | T | C | MB43 | HRSMB26<br>_0090 | RNA ligase | nonsynonymous | I56523T |
| MB43 | C | A | MB43 | HRSMB26<br>_0096 | DNA<br>topoisomerase<br>II | nonsynonymous | R62380S |
| MB43 | A | C | MB43 | HRSMB26<br>_0140 | baseplate tail<br>tube cap | nonsynonymous | I96959M |
| MB43 | A | C | MB43 | HRSMB26<br>_0154 | baseplate<br>wedge subunit | nonsynonymous | K109457T |
| MB43 | A | C | MB43 | HRSMB26<br>_0155 | baseplate<br>wedge subunit | nonsynonymous | I112959L |

Gene presence and SNP differences may contribute to changes in these phages’ local VI phenotype (Table 1, Table S10, Table S11). For instance, the presence of hypothetical protein HRSMB16_0091, adjacent to the holin gene, and unique SNP differences may contribute to the increase in local virulence in MB16 at MOI 0.01. The unique SNP differences and presence of unique genes HRSMB26_0102, HRSMB26_0103 and HRSMB26_0119 and absence genes HRSMB15_0100 and HRSMB15_0116 from MB26 could contribute to the significant increase in local VI at MOI 0.0001. The absence of genes HRSMB16_0023 and HRSMB16_0024, that are situated adjacent to tRNAs in the genome, could contribute to the significant decrease in local VI of MB15 at MOI 0.0001. Future phenotypic work, including a full VI would provide more insight. MB43 had the most unique nonsynonymous SNP differences (Table 1), including changes in three genes related to base plate proteins, we predict that these changes influence the reduced host range we observed in this phage. Future work focusing on the application of these phages for therapeutic use would benefit from further phage isolation to increase host range and optimising phage formulation and delivery, perhaps using similar a strategy as current antibiotic treatment applications (e.g. topical or liquid injectable).

## Conclusion

This study describes the isolation and characterisation of the four novel *Moraxella bovis* phages MB15, MB16, MB26 and MB43. We assessed their phenotypic traits, host range specificity and compared their genomes. These phages are genetically similar, with 8 gene changes and 35 SNP differences, they represent a new genus and species and are taxonomically classified with taxMyPhage as within the family *Straboviridae*, named “Nasusvirus vacca*”*. The phages infected 21 of 41 clinical isolates, notably MB43 had a reduced host range, just infecting 16 of the 41 strains tested. MB16 has the highest local VI at MOI 0.01 and MB26 had the highest local VI at MOI 0.0001. This work demonstrates the importance of phenotypic characterisation of genetically similar phages. Phenotypic characterisation, in addition to genomic analysis, is essential to find the best phages for therapy and to better understand phage biology. Our data provides a strong starting point for further research, supporting the therapeutic potential of these phages in future phage therapy applications for *Moraxella* infectious diseases.

## Acknowledgments

This research was funded by Carus Animal Health Ltd. HRS and MW were supported by; BMCS, HLP, DKK, AR are employees of and GF and RO work for Carus Animal Health Ltd. AM was supported by MRC-CLIMB (MR/T030062/1). AMT is funded by BBSRC (BB/Y51374X/1). AK was supported by MRC AIM Grant Ref: MR/W007002/1.

We thank Orhan Sahin at Iowa State University, for the provision of the *M. bovis* clinical isolates. Thanks to Natalie Allcock and Mischa Haria for their assistance with processing and capturing the transmission electron microscopy images at the University of Leicester Core Biotechnology Services Electron Microscopy Facility. Thanks to Arezoo Pedramfar for assistance with phage stock aliquots and Slawomir Michniewski for experimental and genome submission advice. This research used the ALICE High Performance Computing facility at the University of Leicester.

HRS: Conceptualization; Data curation; Formal analysis; Investigation; Methodology; Visualization; Writing – original draft. MW: Formal analysis; Investigation; Methodology; Visualization. TJ, NIU, AK, AMT: Formal analysis; Methodology. DKK, AR, HLP, BMCS: Formal analysis; Investigation; Methodology. GF: Conceptualization; Investigation; Resources. RO: Conceptualization; Resources. ADM: Formal analysis; Methodology; Validation. MRJC: Conceptualization; Methodology; Resources; Supervision; Validation. All authors: Writing – review & editing.

TJ contributed to One-Step and adsorption assays; NIU, AK and AT contributed to LVI assays; DKK, AR, HLP and BMCS contributed to host range analysis; GF collected swab samples.

