## Supplementary Information for "Characterisation of novel bacteriophages against the cattle pathogen *Moraxella bovis*"

Table S1: *M. bovis* clinical isolates used in this study.

| Study ID | IOWA ID | Year | specimen | Species | Owner State | Site State | Breed Name |
|---|---|---|---|---|---|---|---|
| ISO6 | 7 | 2017 | Eye swab | *Moraxella bovis* | IA | IA | Crossbreed |
| ISO7 | 8 | 2017 | Eye swab | *Moraxella bovis* | IA | IA | Crossbreed |
| ISO8 | 10 | 2017 | Eye swab | *Moraxella bovis* | IA | IA | Crossbreed |
| ISO9 | 11 | 2017 | Eye swab | *Moraxella bovis* | IA | IA | Unknown |
| ISO10 | 12 | 2017 | Eye swab | *Moraxella bovis* | IA | IA | Other |
| ISO11 | 15 | 2017 | Eye swab | *Moraxella bovis* | IA | IA | Salers |
| ISO12 | 19 | 2017 | Eye swab | *Moraxella bovis* | UNKNOWN | UNKNOWN | Crossbreed |
| ISO13 | 23 | 2017 | Eye swab | *Moraxella bovis* | NC | NC | Holstein |
| ISO14 | 28 | 2017 | Eye swab | *Moraxella bovis* | IA | IA | Angus |
| ISO15 | 30 | 2017 | Eye swab | *Moraxella bovis* | NE | NE | Unknown |
| ISO16 | 34 | 2018 | Eye swab | *Moraxella bovis* | SD | SD | Unknown |
| ISO18 | 47 | 2018 | Eye swab | *Moraxella bovis* | IA | IA | Angus |
| ISO19 | 50 | 2018 | Eye swab | *Moraxella bovis* | IA | IA | Unknown |
| ISO20 | 51 | 2018 | Eye | *Moraxella bovis* | WV | WV | Angus |
| ISO21 | 53 | 2018 | Eye swab | *Moraxella bovis* | IA | IA | Unknown |
| ISO22 | 54 | 2018 | Eye swab | *Moraxella bovis* | MN | UNKNOWN | Holstein |
| ISO25 | 69 | 2018 | Eye swab | *Moraxella bovis* | IA | IA | Angus |
| ISO26 | 75 | 2018 | Eye | *Moraxella bovis* | IA | IA | Angus |
| ISO27 | 79 | 2019 | Eye swab | *Moraxella bovis* | IL | IL | Crossbreed |
| ISO28 | 80 | 2019 | Eye swab | *Moraxella bovis* | ID | ID | Crossbreed |
| ISO29 | 83 | 2019 | Eye | *Moraxella bovis* | IA | IA | Holstein |
| ISO30 | 85 | 2019 | Eye swab | *Moraxella bovis* | IA | IA | Holstein |
| ISO32 | 94 | 2019 | Eye swab | *Moraxella bovis* | IA | UNKNOWN | Unknown |

| | | | | | | | |
|---|---|---|---|---|---|---|---|
| ISO33 | 97 | 2019 | Eye swab | *Moraxella bovis* | IA | IA | Holstein |
| ISO34 | 101 | 2019 | Eye swab | *Moraxella bovis* | IA | IA | Crossbreed |
| ISO35 | 103 | 2019 | Corneal swab | *Moraxella bovis* | IA | IA | Holstein |
| ISO36 | 105 | 2019 | Eye swab | *Moraxella bovis* | IA | IA | Crossbreed |
| ISO37 | 112 | 2019 | Eye swab | *Moraxella bovis* | IL | IL | Unknown |
| ISO40 | 120 | 2020 | Eye swab | *Moraxella bovis* | IA | MO | Crossbreed |
| ISO41 | 124 | 2020 | Eye swab | *Moraxella bovis* | IA | IA | Beef |
| ISO42 | 126 | 2020 | Eye swab | *Moraxella bovis* | IA | IA | Crossbreed |
| ISO43 | 129 | 2020 | Eye swab | *Moraxella bovis* | UNKNOWN | UNKNOWN | Crossbreed |
| ISO45 | 131 | 2020 | Eye swab | *Moraxella bovis* | IL | IL | Crossbreed |
| ISO46 | 140 | 2020 | Eye swab | *Moraxella bovis* | MN | MN | Beef |
| ISO47 | 141 | 2020 | Eye swab | *Moraxella bovis* | IA | IA | Angus |
| ISO48 | 142 | 2020 | Ocular swab | *Moraxella bovis* | IA | IA | Unknown |
| ISO50 | 148 | 2020 | Eye swab | *Moraxella bovis* | IA | IA | Unknown |
| ISO51 | 150 | 2021 | Eye swab | *Moraxella bovis* | UNKNOWN | UNKNOWN | Crossbreed |
| ISO52 | 157 | 2021 | Eye swab | *Moraxella bovis* | IA | IA | Holstein |
| ISO55 | 165 | 2021 | Ocular swab | *Moraxella bovis* | OK | OK | Crossbreed |
| ISO58 | 187 | 2022 | Eye swab | *Moraxella bovis* | IA | IA | Crossbreed |

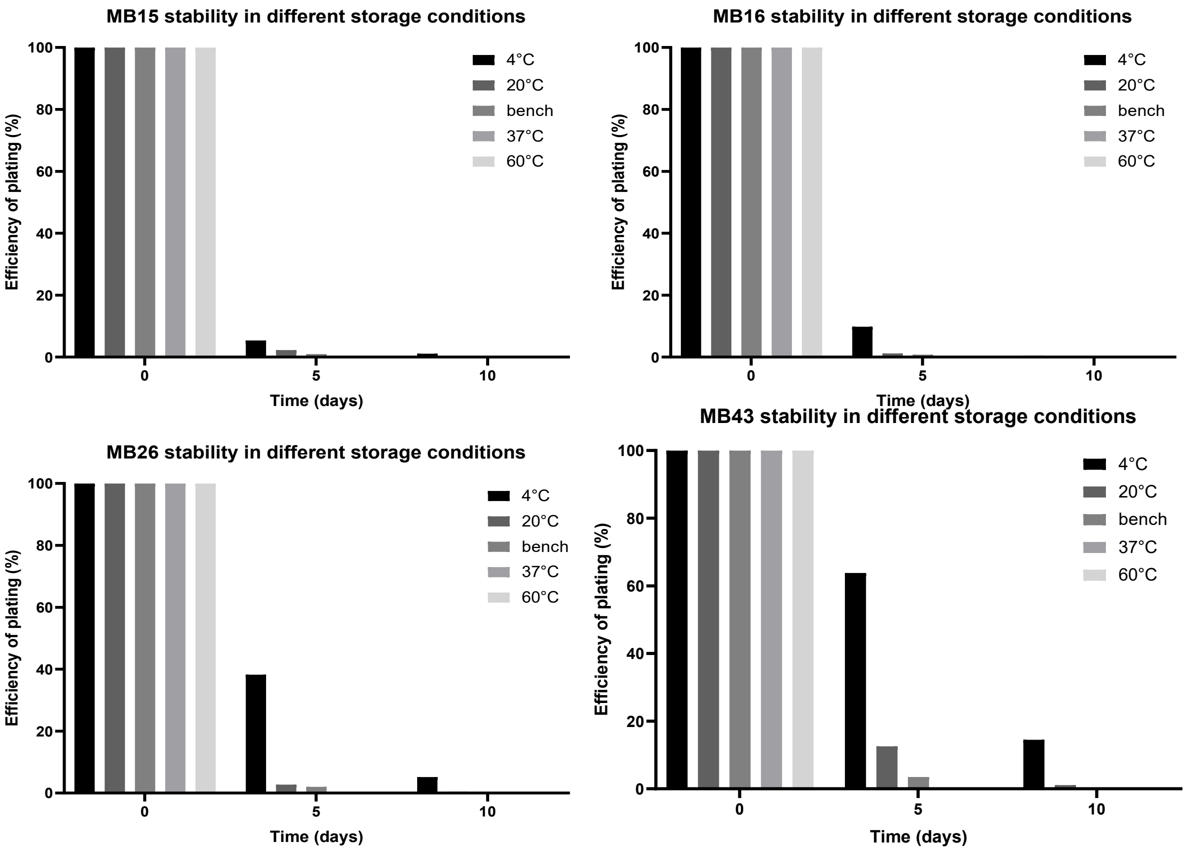


Figure S1: Assessment of phage stability in different temperatures. Phage stocks (MB15, MB16, MB26 and MB43) were aliquoted and stored at different temperatures. Phage titre was periodically measured using the agar overlay spot assay. Phage titre was compared to the starting PFU/mL to calculate the efficiency of plating (%). Phage titres decline rapidly in standard 4 °C lab stocks. No phages were recovered after 5 days storage at 37 °C or 60 °C.

Table S2: Genome information of phages MB15, MB16, MB26 and MB43

| Phage | MB15 | MB16 | MB26 | MB43 |
|---|---|---|---|---|
| Genome length (bp) | 123307 | 123307 | 123294 | 123307 |
| Number of Genes | 161 | 164 | 164 | 163 |
| GC content (%) | 36.0474 | 36.0507 | 36.0472 | 36.0474 |
| Sample number | ERS25317165 | ERS25317166 | ERS26929364 | ERS25317164 |
| Taxonomy ID | 3434924 | 3434925 | 3465768 | 3434927 |
| BioProject | PRJEB81897 | PRJEB81897 | PRJEB81897 | PRJEB81897 |
| Accession Number | OZ335332 | OZ335334 | OZ335335 | OZ335333 |
| Sequencing depth (x) | 5854 | 5811 | 6963 | 6659 |

Table S3: TEM measurements of phage components. Three virions and components were measured, and the mean and standard error (SE) are shown.

| | Tail length (nm) | Head width (nm) | Head length (nm) |
|---|---|---|---|
| MB15 | 125 ± 1.2 | 86 ± 2.3 | 116 ± 2.2 |
| MB16 | 118 ± 1.7 | 82 ± 2.7 | 118 ± 2.0 |
| MB26 | 113 ± 3.0 | 84 ± 1.6 | 118 ± 1.0 |
| MB43 | 114 ± 1.8 | 83 ± 2.3 | 119 ± 1.6 |

Table S4: Local virulence index of phages MB15, MB16, MB26 and MB43 infecting *M. bovis* NCTC 9426. The mean and standard deviation (SD) of three biological repeats are shown.

| | MOI=0.01 | MOI=0.001 | MOI=0.0001 |
|---|---|---|---|
| MB15 | 0.71 ± 0.01 | 0.57 ± 0.01 | 0.42 ± 0.02 |
| MB16 | 0.77 ± 0.02 | 0.62 ± 0.04 | 0.48 ± 0.02 |
| MB26 | 0.72 ± 0.005 | 0.61 ± 0.02 | 0.53 ± 0.02 |
| MB43 | 0.72 ± 0.01 | 0.59 ± 0.04 | 0.47 ± 0.004 |

Table S5. ANOVA of local VI values from Table S7

| | Df | Sum Sq | Mean Sq | F value | Pr(>F) |
|---|---|---|---|---|---|
| phage | 3 | 0.017672 | 0.005891 | 13.73576 | 2.02E-05 |
| MOI | 2 | 0.392839 | 0.196419 | 458.0051 | 7.67E-20 |
| phage:MOI | 6 | 0.010916 | 0.001819 | 4.24211 | 0.004752 |
| Residuals | 24 | 0.010293 | 0.000429 | NA | NA |

Table S6. Tukey-adjusted post-hoc tests of Table S8 ANOVA comparing MOI by phage

| contrast | phage | estimate | SE | df | t.ratio | p.value |
|---|---|---|---|---|---|---|
| MOI0.0001 - MOI0.001 | MB15 | -0.15206 | 0.016909 | 24 | -8.99312 | 1.10E-08 |
| MOI0.0001 - MOI0.01 | MB15 | -0.29347 | 0.016909 | 24 | -17.356 | 3.49E-14 |
| MOI0.001 - MOI0.01 | MB15 | -0.14141 | 0.016909 | 24 | -8.36291 | 4.20E-08 |
| MOI0.0001 - MOI0.001 | MB16 | -0.13988 | 0.016909 | 24 | -8.27244 | 5.11E-08 |
| MOI0.0001 - MOI0.01 | MB16 | -0.29492 | 0.016909 | 24 | -17.4417 | 3.34E-14 |
| MOI0.001 - MOI0.01 | MB16 | -0.15504 | 0.016909 | 24 | -9.16929 | 7.67E-09 |
| MOI0.0001 - MOI0.001 | MB26 | -0.0824 | 0.016909 | 24 | -4.87298 | 0.000164 |
| MOI0.0001 - MOI0.01 | MB26 | -0.19406 | 0.016909 | 24 | -11.4769 | 9.27E-11 |
| MOI0.001 - MOI0.01 | MB26 | -0.11166 | 0.016909 | 24 | -6.6039 | 2.30E-06 |
| MOI0.0001 - MOI0.001 | MB43 | -0.11116 | 0.016909 | 24 | -6.57424 | 2.47E-06 |
| MOI0.0001 - MOI0.01 | MB43 | -0.24062 | 0.016909 | 24 | -14.2306 | 1.02E-12 |
| MOI0.001 - MOI0.01 | MB43 | -0.12946 | 0.016909 | 24 | -7.65631 | 2.00E-07 |

Table S7. Tukey-adjusted post-hoc tests of Table S8 ANOVA comparing phage by MOI

| contrast | MOI | estimate | SE | df | t.ratio | p.value |
|---|---|---|---|---|---|---|
| MB15 - MB16 | 0.0001 | -0.05803 | 0.016909 | 24 | -3.43197 | 0.010931 |
| MB15 - MB26 | 0.0001 | -0.10806 | 0.016909 | 24 | -6.39064 | 7.47E-06 |
| MB15 - MB43 | 0.0001 | -0.05619 | 0.016909 | 24 | -3.32316 | 0.014112 |
| MB16 - MB26 | 0.0001 | -0.05003 | 0.016909 | 24 | -2.95867 | 0.03236 |
| MB16 - MB43 | 0.0001 | 0.00184 | 0.016909 | 24 | 0.108816 | 0.999525 |
| MB26 - MB43 | 0.0001 | 0.051867 | 0.016909 | 24 | 3.067483 | 0.025374 |
| MB15 - MB16 | 0.001 | -0.04584 | 0.016909 | 24 | -2.71129 | 0.055282 |
| MB15 - MB26 | 0.001 | -0.03839 | 0.016909 | 24 | -2.2705 | 0.133353 |
| MB15 - MB43 | 0.001 | -0.01529 | 0.016909 | 24 | -0.90428 | 0.802687 |
| MB16 - MB26 | 0.001 | 0.007453 | 0.016909 | 24 | 0.440787 | 0.970756 |
| MB16 - MB43 | 0.001 | 0.030554 | 0.016909 | 24 | 1.807011 | 0.294673 |
| MB26 - MB43 | 0.001 | 0.023101 | 0.016909 | 24 | 1.366223 | 0.53171 |
| MB15 - MB16 | 0.01 | -0.05948 | 0.016909 | 24 | -3.51767 | 0.008922 |
| MB15 - MB26 | 0.01 | -0.00865 | 0.016909 | 24 | -0.51149 | 0.955548 |
| MB15 - MB43 | 0.01 | -0.00334 | 0.016909 | 24 | -0.19768 | 0.997187 |
| MB16 - MB26 | 0.01 | 0.050831 | 0.016909 | 24 | 3.006177 | 0.029115 |
| MB16 - MB43 | 0.01 | 0.056137 | 0.016909 | 24 | 3.319991 | 0.014216 |
| MB26 - MB43 | 0.01 | 0.005306 | 0.016909 | 24 | 0.313814 | 0.98902 |

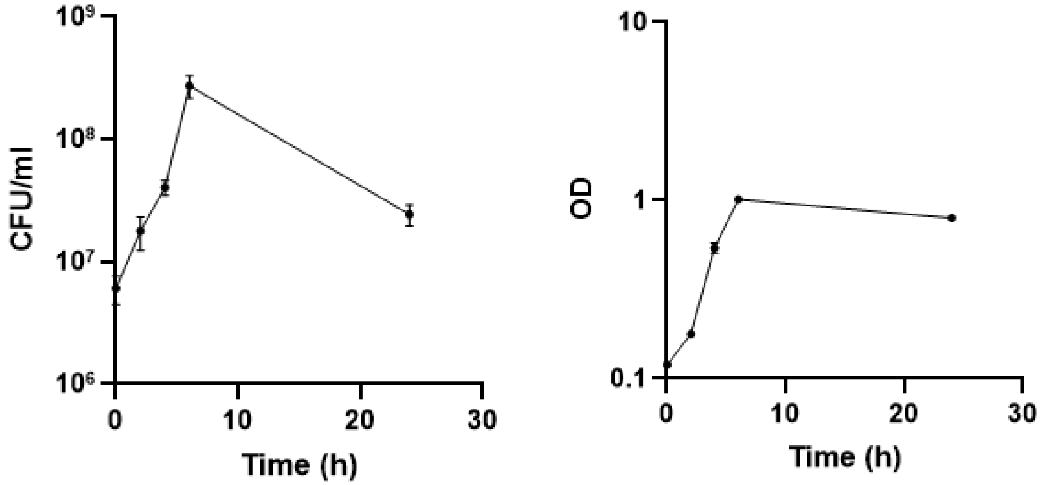


Figure S2: Growth curve of *M. bovis* NCTC 9426. *M. bovis* was grown in BHI+ from a starting optical density (OD 600nm) of 0.1 and incubated at 37 ˚C and 200 rpm for 24 hours. Timepoints were taken at 0 hours, 2 hours, 4 hours 6 hours and 24 hours to monitor CFU/mL (left) and OD (right) over time. Bars represent standard error of the mean, n=3.

Table S8. Phage titres (PFU/mL) data for overnight culture lawns of *M. bovis* isolates susceptible to phage infection.

| *M. bovis* Isolates | Bacteriophage Titres (PFU/mL) | | | |
|---|---|---|---|---|
| | MB15 | MB16 | MB26 | MB43 |
| ISO7 | 1.1E+08 | 1.5E+07 | 8.0E+07 | N/A |
| ISO9 | 3.7E+06 | 9.3E+05 | 9.0E+05 | Visible plaques – Faint- difficult to count |
| ISO10 | 6.8E+07 | 1.4E+08 | 1.6E+08 | 3.5E+08 |
| ISO12 | 1.53E+06 | 3.33E+06 | 2.00E+06 | 1.60E+05 |
| ISO13 | 1.0E+06 | 6.0E+05 | 1.3E+06 | 5.33E+06 |
| ISO15 | 1.5E+08 | 1.8E+08 | 1.5E+08 | 3.7E+07 |
| ISO16 | 2.73E+07 | 8.00E+07 | 8.00E+07 | 4.00E+07 |
| ISO18 | 4.7E+07 | 7.3E+07 | 9.3E+07 | 1.1E+08 |
| ISO20 | 1.6E+08 | 2.2E+08 | 1.0E+07 | Visible plaques – Faint- difficult to count |
| ISO21 | 1.40E+06 | 1.46E+06 | 1.66E+06 | 1.46E+05 |
| ISO22 | 2.1E+08 | 1.5E+08 | 1.9E+08 | 1.0E+08 |
| ISO25 | 1.7E+08 | 1.6E+08 | 1.3E+08 | 1.3E+07 |
| ISO29 | 3.6E+08 | 4.9E+08 | 2.9E+08 | *3.0E+06 |
| ISO30 | 5.0E+08 | 5.7E+08 | 3.7E+08 | 4.7E+07 |
| ISO36 | 1.2E+08 | 1.3E+07 | 2.1E+07 | N/A |
| ISO37 | 1.0E+07 | 1.3E+07 | 1.6E+07 | 1.3E+07 |
| ISO41 | 7.4E+07 | 2.1E+07 | 4.9E+07 | 1.10E+05 |
| ISO46 | 2.3E+08 | 2.7E+08 | 1.9E+08 | N/A |
| ISO48 | 3.8E+07 | 2.2E+06 | 6.9E+06 | *2.86E+07 |
| ISO50 | 2.60E+05 | 1.00E+06 | 4.40E+05 | 7.00E+03 |
| ISO52 | 1.0E+07 | 8.7E+05 | 8.0E+05 | 6.2E+06 |
| NCTC 9426 | 6.66E+06 | 2.60E+07 | 1.06E+07 | 6.66E+06 |

*N=3, N/A: Resistant*, * plaques countable on one occasion

Table S9. Phage titres (PFU/mL) data for mid-exponential culture lawns of *M. bovis* isolates.

| *M. bovis* Isolates | Bacteriophage Titres (PFU/mL) | | | |
|---|---|---|---|---|
| | MB15 | MB16 | MB26 | MB43 |
| ISO6 | Visible plaques, Faint difficult to count | Visible plaques, Faint difficult to count | N/A | N/A |
| ISO7 | N/A | N/A | N/A | N/A |
| ISO9 | N/A | N/A | N/A | N/A |
| ISO10 | 5.40E+06 | 5.90E+06 | 5.07E+06 | 3.07E+06 |
| ISO11 | 2.00E+05 | 6.00E+05 | 2.10E+05 | 1.60E+05 |
| ISO12 | N/A | N/A | N/A | N/A |
| ISO13 | 1.20E+06 | 7.00E+05 | 4.70E+05 | 9.50E+05 |
| ISO15 | 3.80E+06 | 3.00E+06 | 2.00E+05 | 9.50E+05 |
| ISO16 | 3.80E+07 | 3.93E+07 | 1.30E+07 | 7.00E+06 |
| ISO18 | 3.20E+05 | 2.13E+05 | 3.47E+05 | 5.53E+05 |
| ISO20 | 1.35E+06 | 1.47E+06 | 4.40E+06 | 1.90E+06 |
| ISO21 | N/A, Faint lawn, difficult to count | N/A, Faint lawn, difficult to count | N/A, Faint lawn, difficult to count | N/A, Faint lawn, difficult to count |
| ISO22 | 3.13E+05 | 1.87E+05 | 3.20E+05 | 4.93E+05 |
| ISO25 | 4.20E+05 | 3.90E+05 | 1.40E+05 | 2.67E+05 |
| ISO26 | 4.00E+05 | 1.60E+05 | 3.00E+05 | 3.20E+05 |
| ISO28 | Visible plaques, Faint difficult to count | Visible plaques, Faint difficult to count | Visible plaques, Faint difficult to count | Visible plaques, Faint difficult to count |
| ISO29 | 7.00E+05 | 1.00E+06 | 1.60E+05 | 3.33E+05 |
| ISO30 | Visible plaques, Faint difficult to count | Visible plaques, Faint difficult to count | Visible plaques, Faint difficult to count | Visible plaques, Faint difficult to count |
| ISO36 | 3.30E+05 | 4.20E+06 | 1.60E+06 | 8.00E+05 |
| ISO37 | 3.60E+04 | 4.10E+04 | 3.90E+04 | N/A |
| ISO41 | N/A | N/A | 2800 | 1200 |

| | | | | |
|---|---|---|---|---|
| ISO46 | 16000 | 27000 | 28670 | 12000 |
| ISO48 | N/A | N/A | N/A | N/A |
| ISO50 | N/A | N/A | N/A | N/A |
| ISO52 | N/A | N/A | N/A | N/A |
| NCTC 9426 | 9942857 | 1E+07 | 9447143 | 3733333 |

*N=3, N/A: Resistant*, NCTC 9426 N=7

Figure S3: Pharokka plot of *Moraxella* phage MB26. Colours denote gene function.

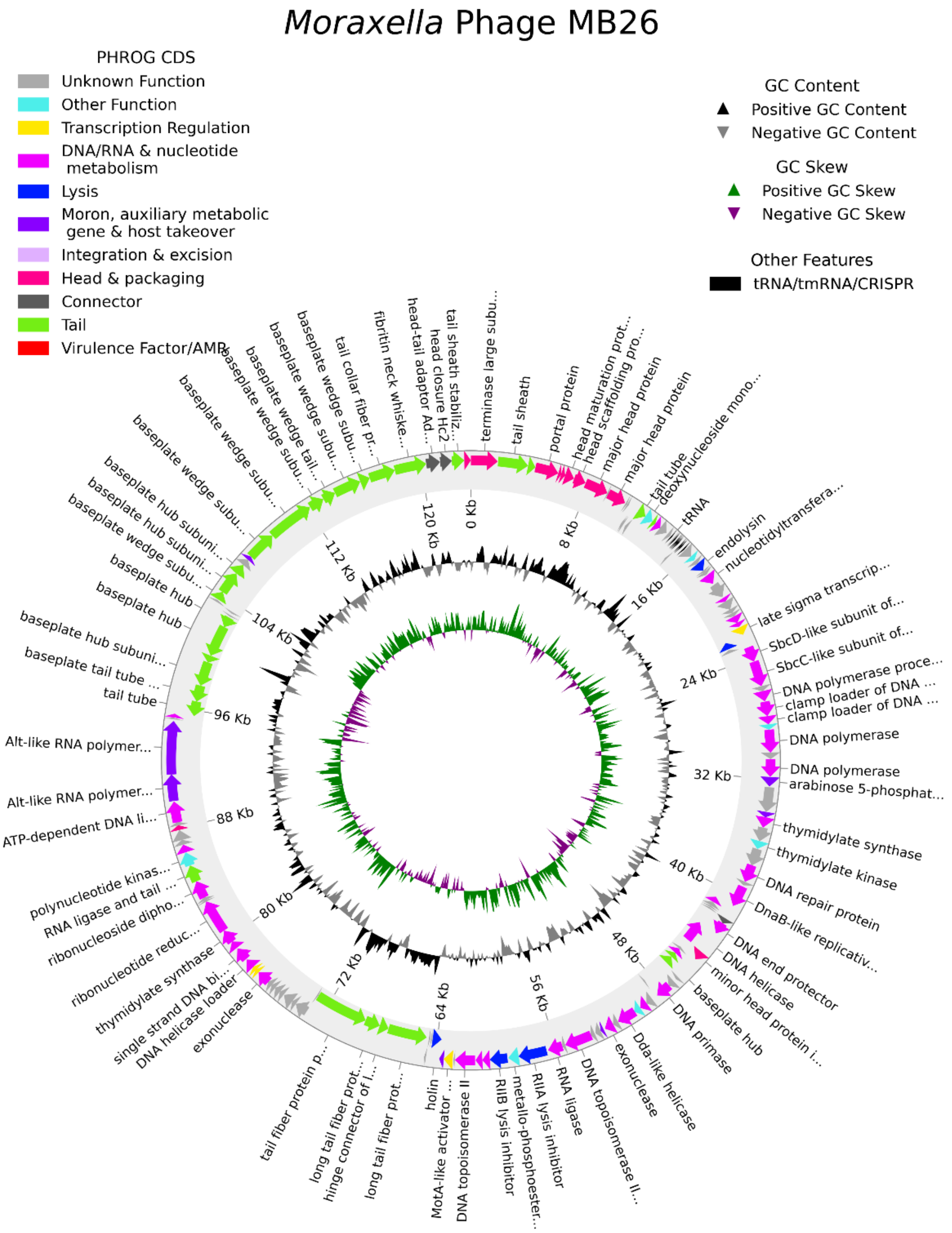

Table S10: Accessory genes in MB15, MB16, MB26 and MB43.

| Gene | MB15 | MB16 | MB26 | MB43 | Annotation | Locus tag | presence _count |
|---|---|---|---|---|---|---|---|
| cluster _160 | 1 | 1 | 0 | 1 | hypothetical protein | HRSMB15_0100, HRSMB16_0103, HRSMB43_0102 | 3 |
| cluster _161 | 1 | 1 | 0 | 1 | DNA helicase loader | HRSMB15_0116, HRSMB16_0119, HRSMB43_0118 | 3 |
| cluster _162 | 0 | 1 | 1 | 1 | hypothetical protein | HRSMB16_0023, HRSMB26_0023, HRSMB43_0023 | 3 |
| cluster _163 | 0 | 1 | 1 | 1 | hypothetical protein | HRSMB16_0024, HRSMB26_0024, HRSMB43_0024 | 3 |

Table S11: Unique genes in MB16 and MB26.

| Gene | MB15 | MB16 | MB26 | MB43 | Annotation | Locus tag | presence _count |
|---|---|---|---|---|---|---|---|
| cluster_ 164 | 0 | 1 | 0 | 0 | hypothetical protein | HRSMB16_0091 | 1 |
| cluster_ 165 | 0 | 0 | 1 | 0 | hypothetical protein | HRSMB26_0102 | 1 |
| cluster_ 166 | 0 | 0 | 1 | 0 | hypothetical protein | HRSMB26_0103 | 1 |
| cluster_ 167 | 0 | 0 | 1 | 0 | DNA helicase loader | HRSMB26_0119 | 1 |

Table S12: All SNP and indel differences between MB15 (ref) and MB16.

| AA change | Ref NT | Alt NT | GeneID | GeneName | Change |
|---|---|---|---|---|---|
| - | T | G | - | - | intergenic |
| - | A | G | - | - | intergenic |
| - | G | T | - | - | intergenic |
| - | A | G | - | - | intergenic |
| - | A | G | - | - | intergenic |
| L56534V | T | G | HRSMB15_0088 | RNA ligase | nonsynonymous |
| G94353V | G | T | HRSMB15_0133 | Alt-like RNA polymerase ADP-ribosyltransferase | nonsynonymous |
| L94358F | C | T | HRSMB15_0133 | Alt-like RNA polymerase ADP-ribosyltransferase | nonsynonymous |
| L94360L | T | G | HRSMB15_0133 | Alt-like RNA polymerase ADP-ribosyltransferase | synonymous |
| D94363E | T | G | HRSMB15_0133 | Alt-like RNA polymerase ADP-ribosyltransferase | nonsynonymous |
| S94369S | T | G | HRSMB15_0133 | Alt-like RNA polymerase ADP-ribosyltransferase | synonymous |
| F103688F | C | T | HRSMB15_0145 | baseplate wedge subunit | synonymous |
| I109460L | A | C | HRSMB15_0151 | baseplate wedge subunit | nonsynonymous |
| N109471K | C | A | HRSMB15_0151 | baseplate wedge subunit | nonsynonymous |

Table S13: All SNP and indel differences between MB15 (ref) and MB26.

| AA change | Ref NT | Alt NT | GeneID | GeneName | Change |
|---|---|---|---|---|---|
| - | T | G | - | - | intergenic |
| - | A | G | - | - | intergenic |
| - | G | T | - | - | intergenic |
| - | T | G | - | - | intergenic |
| F56533L | T | G | HRSMB15_0088 | RNA ligase | nonsynonymous |
| V62372G | T | G | HRSMB15_0094 | DNA topoisomerase II | nonsynonymous |
| - | G | . | HRSMB15_0101 | long tail fiber protein distal subunit | Indel n=13 |
| M66636I | C | A | HRSMB15_0101 | long tail fiber protein distal subunit | nonsynonymous |
| Y94350S | A | C | HRSMB15_0133 | Alt-like RNA polymerase ADP-ribosyltransferase | nonsynonymous |
| L94358F | C | T | HRSMB15_0133 | Alt-like RNA polymerase ADP-ribosyltransferase | nonsynonymous |
| F103688F | C | T | HRSMB15_0145 | baseplate wedge subunit | synonymous |
| N109471K | C | A | HRSMB15_0151 | baseplate wedge subunit | nonsynonymous |
| F112964C | T | G | HRSMB15_0152 | baseplate wedge subunit | nonsynonymous |

Table S14: All SNP and indel differences between MB15 (ref) and MB43.

| AA change | Ref NT | Alt NT | GeneID | GeneName | Change |
|---|---|---|---|---|---|
| - | A | G | - | - | intergenic |
| - | G | T | - | - | intergenic |
| - | G | T | - | - | intergenic |
| I56523T | T | C | HRSMB15_0088 | RNA ligase | nonsynonymous |
| R62380S | C | A | HRSMB15_0094 | DNA topoisomerase II | nonsynonymous |
| K66627K | T | C | HRSMB15_0101 | long tail fiber protein distal subunit | synonymous |
| M66636I | C | A | HRSMB15_0101 | long tail fiber protein distal subunit | nonsynonymous |
| Y94350S | A | C | HRSMB15_0133 | Alt-like RNA polymerase ADP-ribosyltransferase | nonsynonymous |
| L94358F | C | T | HRSMB15_0133 | Alt-like RNA polymerase ADP-ribosyltransferase | nonsynonymous |
| I96972M | A | C | HRSMB15_0137 | baseplate tail tube cap | nonsynonymous |
| F103688F | C | T | HRSMB15_0145 | baseplate wedge subunit | synonymous |
| N109470T | A | C | HRSMB15_0151 | baseplate wedge subunit | nonsynonymous |
| N109471K | C | A | HRSMB15_0151 | baseplate wedge subunit | nonsynonymous |
| I112972L | A | C | HRSMB15_0152 | baseplate wedge subunit | nonsynonymous |

Table S15: All SNP and indel differences between MB16 (ref) and MB26.

| AA change | Ref NT | Alt NT | GeneID | GeneName | Change |
|---|---|---|---|---|---|
| A14614A | G | A | HRSMB16_0023 | hypothetical protein | synonymous |
| S14617R | T | G | HRSMB16_0023 | hypothetical protein | nonsynonymous |
| - | G | A | - | - | intergenic |
| F56533L | T | G | HRSMB16_0090 | RNA ligase | nonsynonymous |
| V56534L | G | T | HRSMB16_0090 | RNA ligase | nonsynonymous |
| V62372G | T | G | HRSMB16_0097 | DNA topoisomerase II | nonsynonymous |
| - | G | . | HRSMB16_0104 | long tail fiber protein distal subunit | Indel n=13 |
| M66636I | C | A | HRSMB16_0104 | long tail fiber protein distal subunit | nonsynonymous |
| Y94350S | A | C | HRSMB16_0136 | Alt-like RNA polymerase ADP-ribosyltransferase | nonsynonymous |
| V94353G | T | G | HRSMB16_0136 | Alt-like RNA polymerase ADP-ribosyltransferase | nonsynonymous |
| L94360F | G | T | HRSMB16_0136 | Alt-like RNA polymerase ADP-ribosyltransferase | nonsynonymous |
| E94363D | G | T | HRSMB16_0136 | Alt-like RNA polymerase ADP-ribosyltransferase | nonsynonymous |
| S94369S | G | T | HRSMB16_0136 | Alt-like RNA polymerase ADP-ribosyltransferase | synonymous |
| L109460I | C | A | HRSMB16_0154 | baseplate wedge subunit | nonsynonymous |
| F112964C | T | G | HRSMB16_0155 | baseplate wedge subunit | nonsynonymous |

Table S16: All SNP and indel differences between MB16 (ref) and MB43.

| AA change | Ref NT | Alt NT | GeneID | GeneName | Change |
|---|---|---|---|---|---|
| - | G | T | - | - | intergenic |
| G14607V | G | T | HRSMB16_0023 | hypothetical protein | nonsynonymous |
| A14614A | G | A | HRSMB16_0023 | hypothetical protein | synonymous |
| - | G | A | - | - | intergenic |
| I56523T | T | C | HRSMB16_0090 | RNA ligase | nonsynonymous |
| V56534L | G | T | HRSMB16_0090 | RNA ligase | nonsynonymous |
| R62380S | C | A | HRSMB16_0097 | DNA topoisomerase II | nonsynonymous |
| K66627K | T | C | HRSMB16_0104 | long tail fiber protein distal subunit | synonymous |
| M66636I | C | A | HRSMB16_0104 | long tail fiber protein distal subunit | nonsynonymous |
| Y94350S | A | C | HRSMB16_0136 | Alt-like RNA polymerase ADP-ribosyltransferase | nonsynonymous |
| V94353G | T | G | HRSMB16_0136 | Alt-like RNA polymerase ADP-ribosyltransferase | nonsynonymous |
| L94360F | G | T | HRSMB16_0136 | Alt-like RNA polymerase ADP-ribosyltransferase | nonsynonymous |
| E94363D | G | T | HRSMB16_0136 | Alt-like RNA polymerase ADP-ribosyltransferase | nonsynonymous |
| S94369S | G | T | HRSMB16_0136 | Alt-like RNA polymerase ADP-ribosyltransferase | synonymous |
| I96972M | A | C | HRSMB16_0140 | baseplate tail tube cap | nonsynonymous |
| L109460I | C | A | HRSMB16_0154 | baseplate wedge subunit | nonsynonymous |
| K109470T | A | C | HRSMB16_0154 | baseplate wedge subunit | nonsynonymous |
| I112972L | A | C | HRSMB16_0155 | baseplate wedge subunit | nonsynonymous |

Table S17: All SNP and indel differences between MB26 (ref) and MB43.

| AA change | Ref NT | Alt NT | GeneID | GeneName | Change |
|---|---|---|---|---|---|
| - | G | T | - | - | intergenic |
| G14607V | G | T | HRSMB26_0023 | hypothetical protein | nonsynonymous |
| R14617S | G | T | HRSMB26_0023 | hypothetical protein | nonsynonymous |
| I56523T | T | C | HRSMB26_0090 | RNA ligase | nonsynonymous |
| L56533F | G | T | HRSMB26_0090 | RNA ligase | nonsynonymous |
| G62372V | G | T | HRSMB26_0096 | DNA topoisomerase II | nonsynonymous |
| R62380S | C | A | HRSMB26_0096 | DNA topoisomerase II | nonsynonymous |
| - | . | G | HRSMB26_0104 | long tail fiber protein distal subunit | Indel n=13 |
| K66614K | T | C | HRSMB26_0104 | long tail fiber protein distal subunit | synonymous |
| I96959M | A | C | HRSMB26_0140 | baseplate tail tube cap | nonsynonymous |
| K109457T | A | C | HRSMB26_0154 | baseplate wedge subunit | nonsynonymous |
| C112951F | G | T | HRSMB26_0155 | baseplate wedge subunit | nonsynonymous |
| I112959L | A | C | HRSMB26_0155 | baseplate wedge subunit | nonsynonymous |
